## Supplementary Figures for "Regulatory networks of KRAB zinc finger genes and transposable elements changed during human brain evolution and disease"

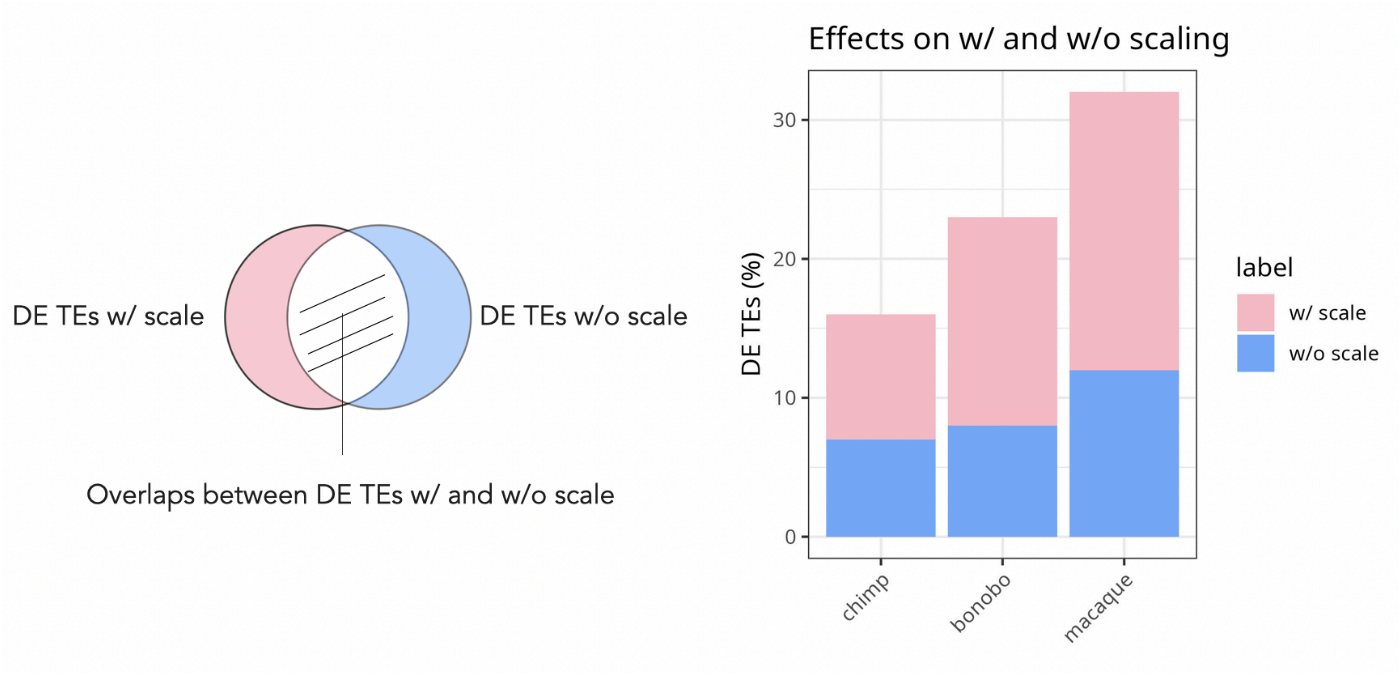


**Figure S1. Comparison of differentially expressed (DE) TEs with and without scaling.** For across species comparison between humans and the NHPs indicated on the x-axis, using expression data from the primary and secondary cortices as an example. The y-axis shows the percentage of TEs that were only called using scaled or non-scaled data (the remainder needed to add up to 100% is the overlap of DE TEs between both methods). To be called DE, the TE needed to show an absolute log2foldchange larger than 1.5 and adjusted *p*-value < 0.05. The impact of scaling is the highest for the most distantly related species, the rhesus macaque, where more than 30% of TEs changed in assignment to being DE or not depending on the applied scaling.

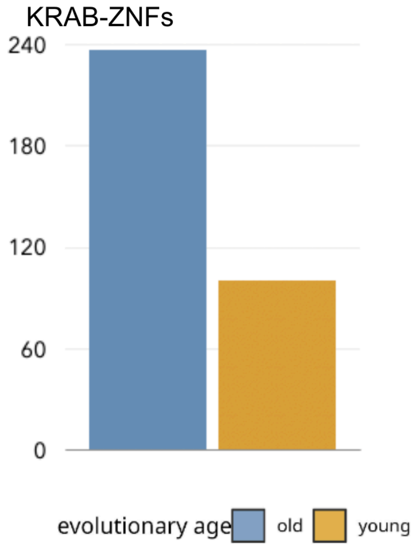


**Figure S2. Distribution of KRAB-ZNFs evolutionary age inference.** The evolutionary age of KRAB-ZNFs was inferred from GenTree (Shao et al., 2019) and primate orthologous annotations (Jovanovic et al., 2021). The evolutionary young group (≤ 44.2 mya) is in orange and the evolutionary old group (> 44.2 mya) is in blue.

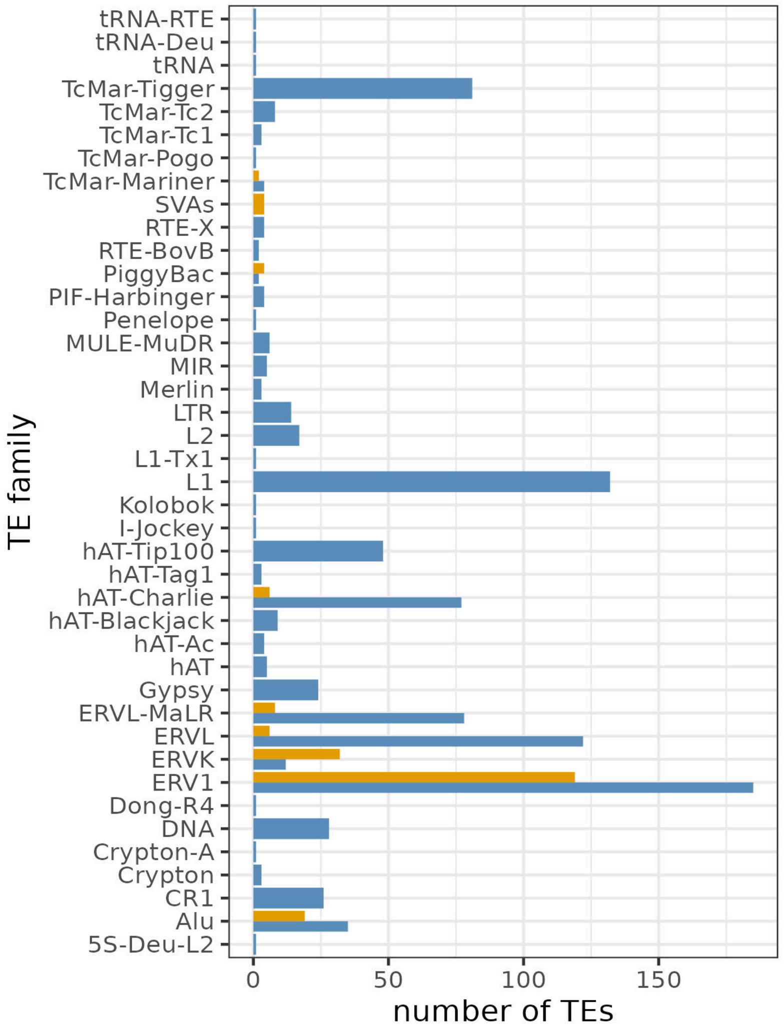


**Figure S3. Distribution of TEs evolutionary age inference.** The age of TEs were estimated using Dfam subfamily species annotation. The evolutionary young group (≤ 44.2 mya) is in orange and the evolutionary old group (> 44.2 mya) is in blue.

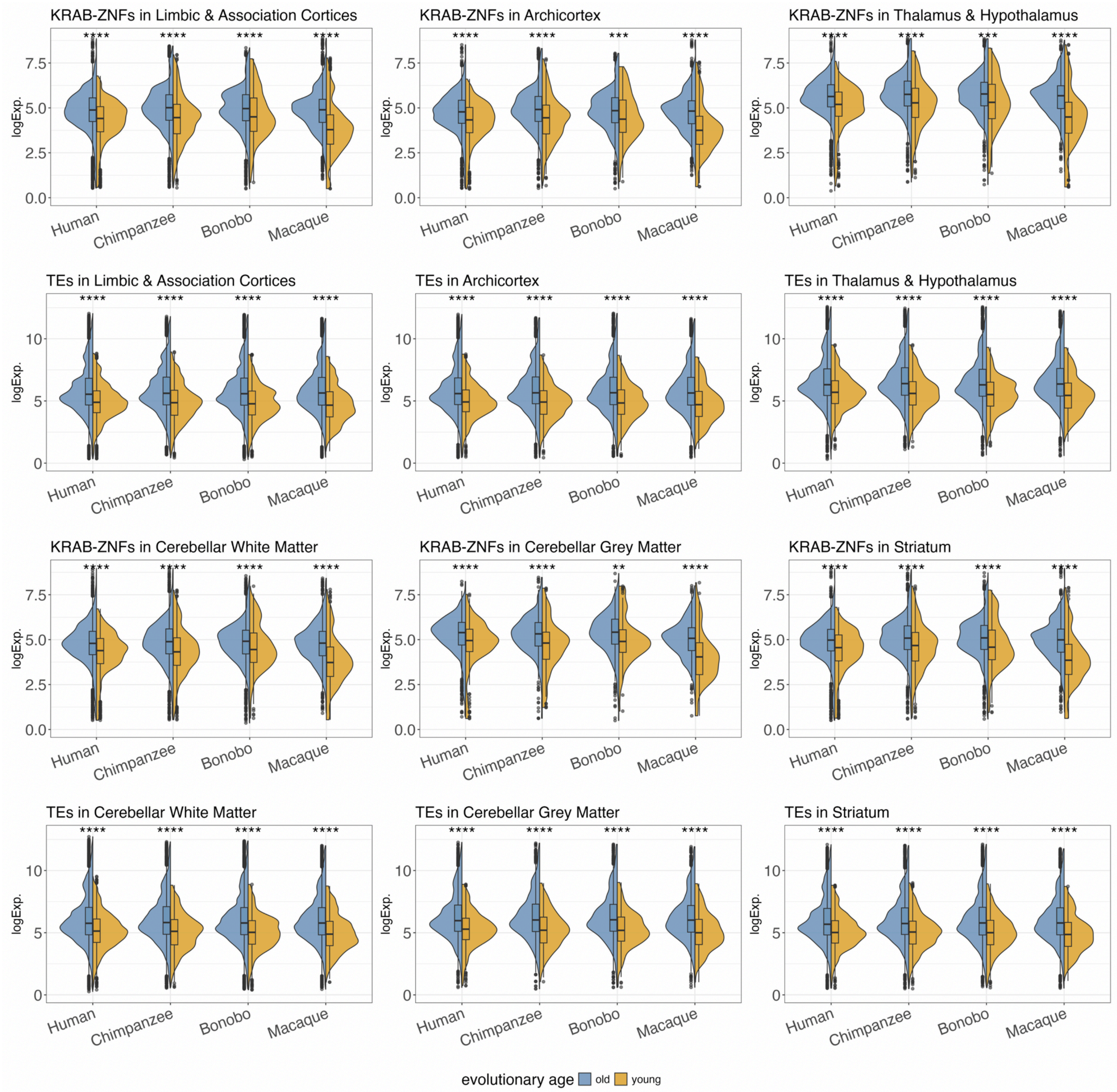


**Figure S4. Expression of KRAB-ZNFs and TEs among brain regions.** Both KRAB-ZNF genes and TEs were grouped in two groups based on their inferred evolutionary age, old (>44.2 mya) and young (≤ 44.2mya). Young KRAB-ZNFs and young TEs have lower expression levels (Wilcoxon Rank Sum Test, $p < 0.05$).

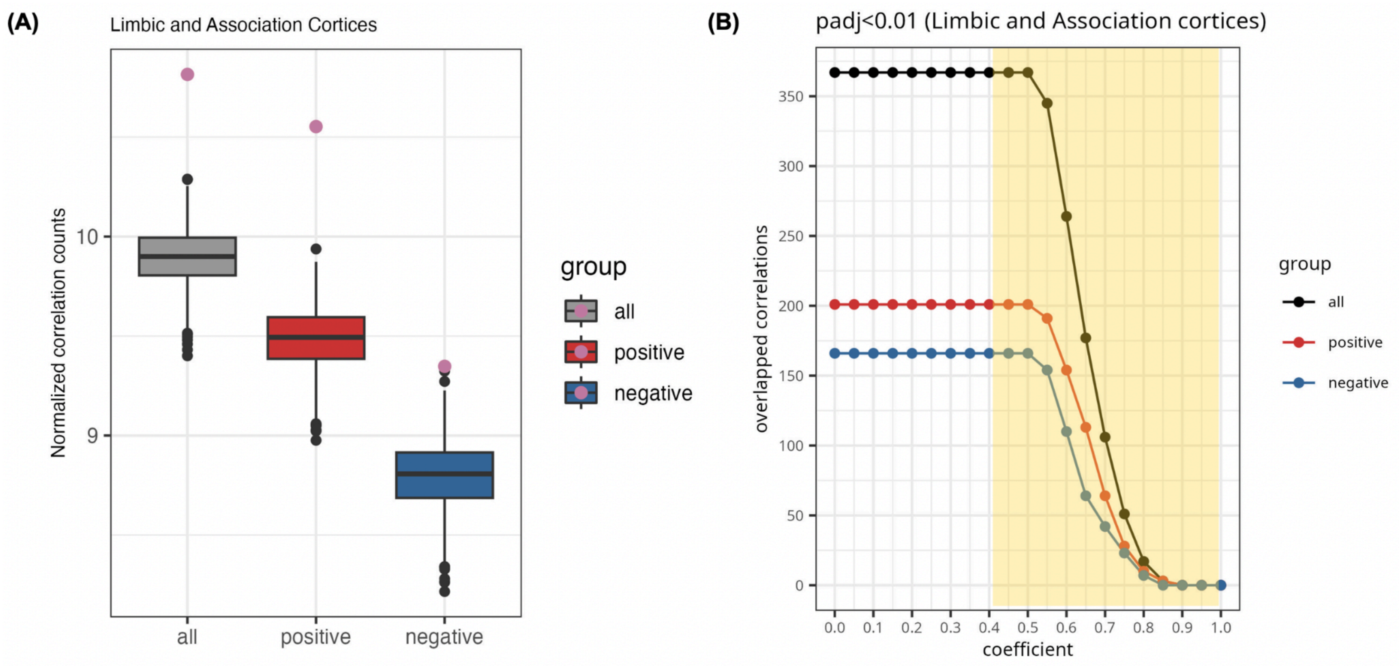


**Figure S5. Distribution of correlation in limbic and association cortices.** This is the same method mentioned in **Figure 3A** and **3B**. **(A)** Higher number of correlations comparing to KRAB-ZNFs (pink dot) and random selected genes ($p$ < 0.001) **(B)** We use a threshold adjusted $p$-value < 0.01 and absolute coefficient larger than 0.4 to select TE:KRAB-ZNF for down-stream analysis.

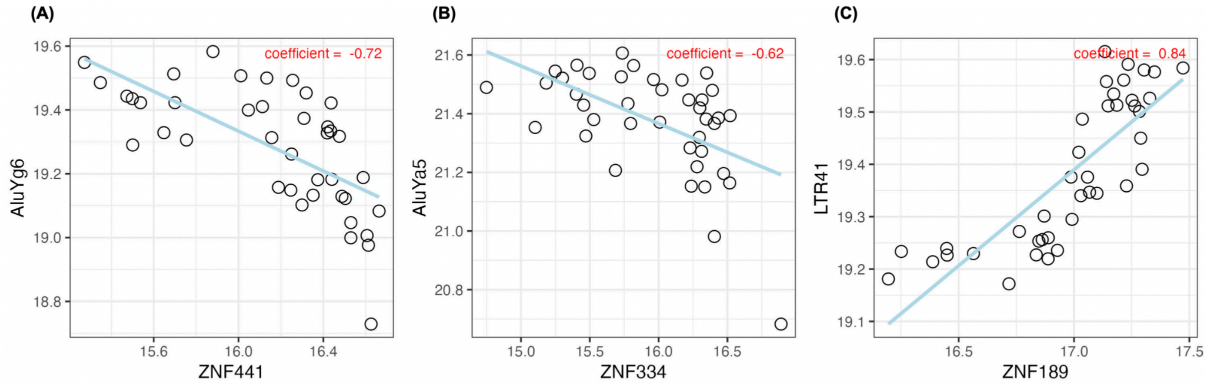


**Figure S6. Example of correlations (TE:KRAB-ZNF).** We define it as a significant one-to-one correlation between TE and KRAB-ZNF based on the absolute coefficient being larger than 0.4 and the adjusted $p$-value being smaller than 0.01. We classify it as a young correlation when at least one of the components (TE or KRAB-ZNF gene) is evolutionary young. **(A)** A negative young example (young TE correlated with young KRAB-ZNF) **(B)** A negative young example (young TE correlated with old KRAB-ZNF) **(C)** An old example (old TE correlated with old KRAB-ZNF). X-axis is the log expression level of KRAB-ZNF and y-axis is the log expression of TE.

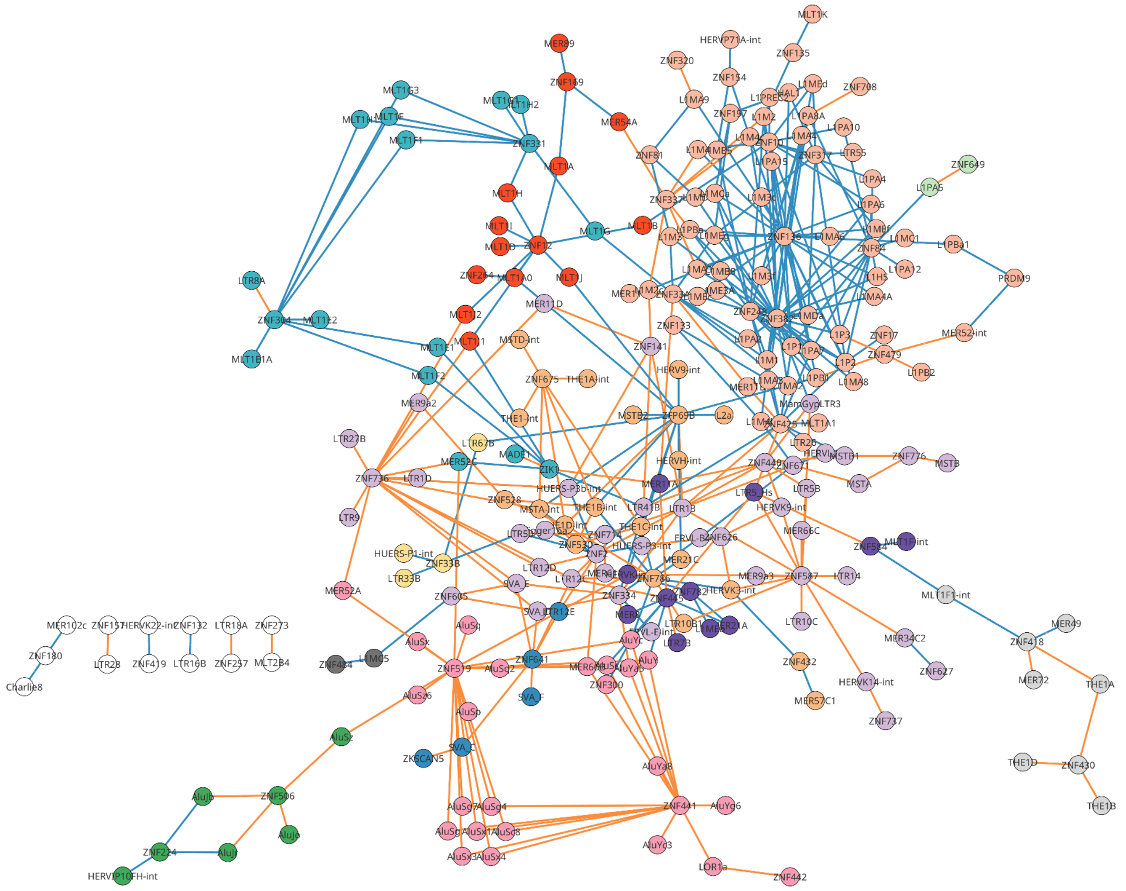

**Figure S7. TE:KRAB-ZNF network in human limbic and association cortices.** Nodes are colored based on 13 different modules. Nodes in white do not belong to any module. Links demonstrate the evolutionary age of the interaction (young link in orange; old link in blue).

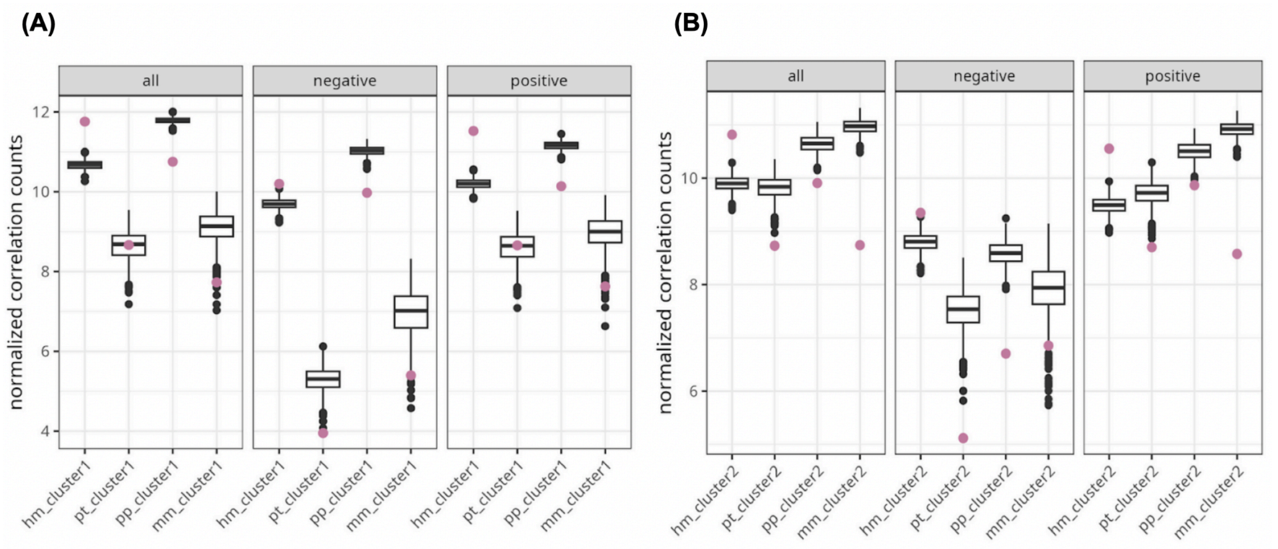


**Figure S8. Comparing KRAB-ZNFs and random selected genes in Primate Brain Data. (A)** primary and secondary cortices (cluster1) and **(B)** limbic and association cortices (cluster2). Violet dots indicating numbers of significant correlations of TE:KRAB-ZNF (adjusted *p* < 0.01). Box plots indicate the distribution of 1000 iterations of random selected genes correlated with TEs (hm: human, pt: chimp, pp: bonobo, mm: macaque, all: positive and negative correlations, negative: negative correlations, positive: positive correlations).

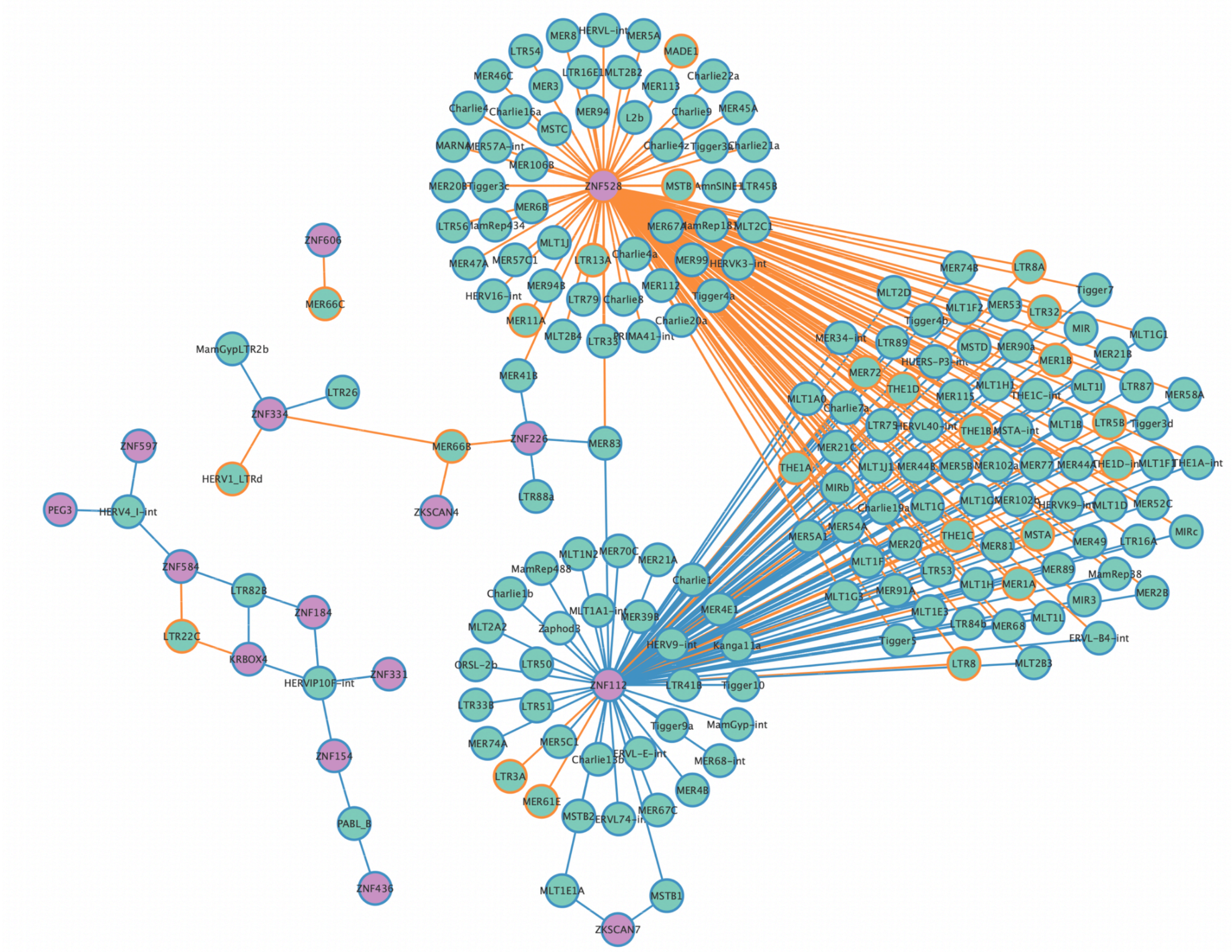


**Figure S9. 276 opposite TE:KRAB-ZNF regulatory network comparing humans to bonobos.** This bipartite network refers to the 276 TE:KRAB-ZNF network mentioned in **Figure 4C**. KRAB-ZNF nodes are in violet and TE nodes are in green. The colors of the border of nodes and the edges represent their evolutionary age. The evolutionary young nodes and edges are in orange and the evolutionary old nodes and edges are in blue.
